# Reducing malaria burden and accelerating elimination with long-lasting systemic insecticides: a modeling study of three potential use cases

**DOI:** 10.1101/615773

**Authors:** Prashanth Selvaraj, Joshua Suresh, Edward A. Wenger, Caitlin A. Bever, Jaline Gerardin

## Abstract

**Background:** While bednets and insecticide spraying have had significant impact on malaria burden in many endemic regions, outdoor vector feeding and insecticide resistance may ultimately limit their contribution to elimination and control campaigns. Complementary vector control methods such as endectocides or systemic insecticides are therefore generating much interest. Here we explore the conditions under which long-lasting systemic insecticides would have a substantial impact on transmission and burden.

**Methods:** Hypothetical long-lasting systemic insecticides with effective durations ranging from 14 days to 90 days are simulated using an individual-based mathematical model of malaria transmission. The impact of systemic insecticides when used to complement existing vector control and drug campaigns is evaluated in three settings – a highly seasonal high-transmission setting, a near-elimination setting with seasonal travel to a high-risk area, and a near-elimination setting in southern Africa.

**Results:** At 60% coverage, a single round of long-lasting systemic insecticide with effective duration of at least 60 days, distributed at the start of the season alongside a seasonal malaria chemoprevention campaign in a high-transmission setting, results in further burden reduction of 30-90% depending on the sub-populations targeted. In a near-elimination setting where transmission is sustained by seasonal travel to a high-risk area, targeting high-risk travelers with systemic insecticide with effective duration of at least 30 days can result in likely elimination even if intervention coverage is as low as 50%. In near-elimination settings with robust vector control, the addition of a 14-day systemic insecticide alongside an antimalarial in mass drug administration (MDA) campaigns can decrease the necessary MDA coverage from about 85% to the more easily achievable 65%.

**Conclusions:** While further research into the safety profile of systemic insecticides is necessary before deployment, we find that long-lasting systemic insecticides can play a critical role in reducing burden or eliminating malaria in a range of contexts with different target populations, existing malaria control methods, and transmission intensities.

## Background

Following the renewed global call for eradication, the widespread rollout of insecticide-treated nets (ITNs), and increased usage of artemisinin-based combination therapies for first-line treatment, malaria burden has declined substantially over the last decade [1]. Yet malaria remains recalcitrant in many areas, including high-burden countries where progress has stalled or reversed in the last few years [2] as well as near-elimination areas where interruption of transmission continues to remain out of reach.

In areas where malaria persists despite high usage of ITNs, outdoor exposure can contribute a major share of residual transmission [3]. Insecticide resistance is increasingly widespread, diminishing the community benefits of ITNs and potentially erasing much of the gains made in the last twenty years of malaria control [4,5]. Continued innovation along the path to eradication is needed to address both challenges.

One potential avenue for complementing the existing toolset is the use of systemic insecticides, known as endectocides when they also have antiparasitic properties, to reduce vector populations. Ivermectin has been distributed in mass drug administrations (MDA) for onchocerciasis and lymphatic filariasis [6], and is lethal to mosquitoes upon ingestion during blood feeding on humans or animals [6,7,8]. However, ivermectin has a short half-life of under 4 days in large mammals [10] and 12-36 hours in humans [11]. At the currently approved doses given for treating non-malarial diseases, ivermectin concentrations in human blood can maintain mosquitocidal effects for approximately 48 hours [12], which is predicted to be too short to have much impact on malaria burden [13]. While the low concentrations of ivermectin observed 28 days post-dose continue to have a small impact on mosquito survival [14], the epidemiological relevance is unknown. The duration of ivermectin efficacy can be extended via multiple doses over the course of weeks, but this could prove operationally challenging to deliver at scale. There has been recent interest in developing longer-lasting systemic insecticide formulations for malaria control and potential elimination, including slow release formulations [12] and high-dose ivermectin, which has been observed to reduce mosquito survival 28 days post-treatment in the field [15]. A recent study in Burkina Faso observed a 20% reduction in clinical incidence in an intervention group receiving ivermectin 6 times over 18 weeks at 3 week intervals, with young children, pregnant women, and women nursing newborns excluded from MDA eligibility [16]. Another drug class, isoxazolines, shows promise to maintain mosquitocidal activity up to 50-90 days [17], although safety concerns still need to be addressed [18].

Previous mathematical modeling has suggested that high-dose ivermectin can marginally improve the impact of antimalarial MDAs [15,19], a new long-lasting systemic insecticide with high efficacy lasting 30 days can substantially reduce burden when distributed with seasonal malaria chemoprevention (SMC) [13], and MDA with an isoxazoline lasting 90 days can also reduce burden in high-transmission areas [17]. However, there has not been a systematic exploration of the potential impact of long-lasting systemic insecticides across multiple use cases within the same modeling framework. In this work, we test the ability of generic hypothetical systemic insecticides lasting 14 to 90 days to achieve additional burden reduction in the context of SMC, targeted elimination in the context of human travel to high-risk areas, and local elimination in the context of outdoor-biting vectors or insufficient coverage with traditional vector control interventions. We predict that long-lasting systemic insecticides can complement other interventions to help reduce burden and achieve elimination. Systemic insecticides increase the impact of drug campaigns or alternatively can be used to achieve the same impact at lower, perhaps more feasible, campaign coverage. Unless systemic insecticides are extremely long-lasting, with mosquitocidal activity up to 90 days post dosage, improving their safety profile such that young children and women of childbearing age can safely participate in MDAs is essential.

## Methods

### Simulation framework

All simulations are performed with EMOD v2.15 [20], an agent-based model of *Plasmodium falciparum* malaria transmission that includes vector life cycle dynamics [21], within-host parasite and immune dynamics calibrated to field parasitology [22], and drug pharmacokinetics and pharmacodynamics [23].

Pharmacokinetics of systemic insecticides are modeled as constant efficacy of duration over 14, 30, 60, or 90 days, subsequently referred to as SI-14, SI-30, SI-60, and SI-90 respectively. Vectors feeding on a human with active systemic insecticide have 95% probability of death prior to the next feed, with three days between feeds. We assume systemic insecticides have identical impact on all vector species and no impact of the systemic insecticide on parasite load. All models include surface area dependent biting, where children are less likely to be bitten by vectors due to their smaller size.

We assume no resistance to antimalarials or to systemic insecticide. In each of the scenarios, coverage refers to the target demographic and not the entire population. For example, 50% coverage in a cohort excluding women of childbearing age and children under 5 would refer to 50% of males over 5 and females between the ages of 5 and 12 and over 51 years of age receiving the systemic insecticide.

### Burden reduction scenarios

We simulate a well-mixed village with Sahelian seasonality and high-intensity transmission of annual entomological inoculation rate (EIR) 110 and overall mean annual prevalence of around 90% in the absence of any interventions. The vector population is modeled as *Anopheles gambiae* mosquitoes with 65% anthropophily and 90% indoor biting. Total human population is around 1000 individuals with birth and death rates of 45 per 1000 per year.

Drug campaigns are timed to start on June 28, just prior to the wet season, and the number of clinical cases averted is calculated for the year after the start of the campaign. Clinical cases are defined as malarial fevers of at least 38.5°C occurring at least 14 days since the previous fever. SMC campaigns are carried out with dihydroartemisinin-piperaquine (DP), which has a 30-day period of prophylactic protection. SMC is distributed as 4 rounds separated by 1 month, with independent coverage between rounds, and targeted at children under 5 in standard SMC or children under 10 in expanded SMC. Systemic insecticide MDAs are given once, concurrently with the first round of SMC. We test three systemic insecticide distribution scenarios: where children under 5 and women of childbearing age (12 – 51 years) [24] are ineligible to receive long-lasting systemic insecticides; where children under 5 are ineligible to receive long-lasting systemic insecticides but women of childbearing age are eligible; and where everyone is eligible to receive long-lasting systemic insecticides.

In all scenarios, we measure the relative reduction in total clinical burden (number of clinical malaria cases in all age groups) compared with a standard SMC campaign in children under 5. Coverage of SMC and systemic insecticide MDA are assumed to be equivalent, although the denominators differ. Fifty stochastic realizations are run for each drug campaign and coverage combination. No other interventions are included in the simulations.

### Targeted elimination scenarios

We simulate a transmission ecosystem of two villages that share a high-risk area (Figure 2A) [25]. Transmission is seasonal, peaking in December, and overall annual mean prevalence of any infection is 29% in the absence of interventions. All villagers can travel between the villages, but only high-risk travelers visit the high-risk area. Each village is home to about 280 people and there is no importation from outside the modeled areas. Village vectors are *Anopheles minimus* and vectors in the high-risk area are *Anopheles dirus*, with 40% and 99% outdoor biting respectively. Both vector species are modeled with 50% anthropophily.

**Figure 1.**
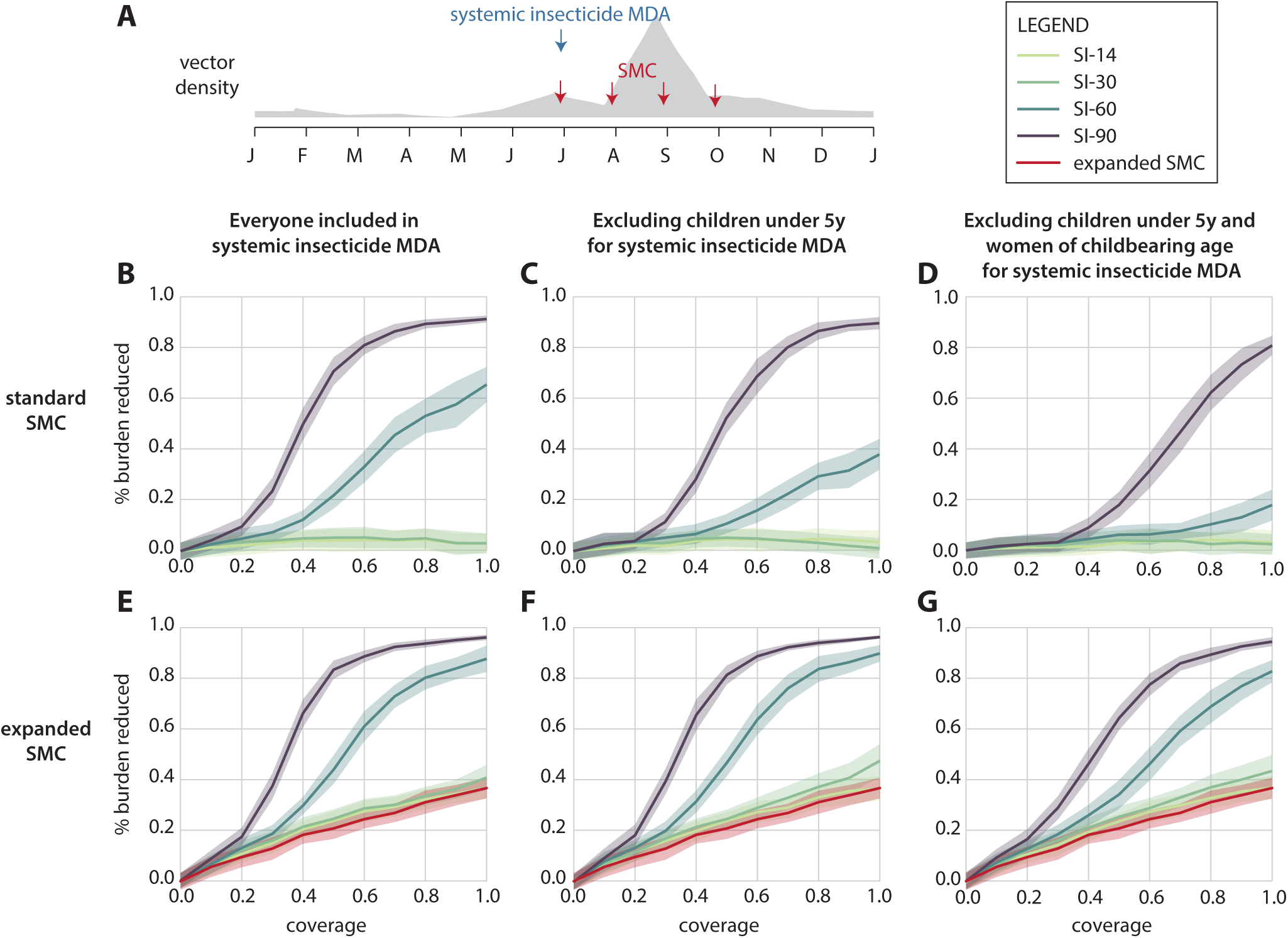
Impact of long-lasting systemic insecticides combined with SMC on burden reduction in the Sahel. (A) Simulated seasonality of adult vectors and timing of SMC and systemic insecticide MDA. (B-G) Impact of systemic insecticides distributed concurrently with the first round of SMC. Coverage is assumed to be the same for both SMC and systemic insecticide MDA. Percentage reduction in total clinical cases is shown in comparison to a standard SMC-only campaign. (B, C, D) Standard SMC campaign in children under 5; (E, F, G) Expanded SMC including all children under 10. (B, E) No restrictions on systemic insecticide eligibility (C, F) Children under 5 ineligible for systemic insecticides (D, G) Children under 5 and women of childbearing age ineligible for systemic insecticides.

**Figure 2.**
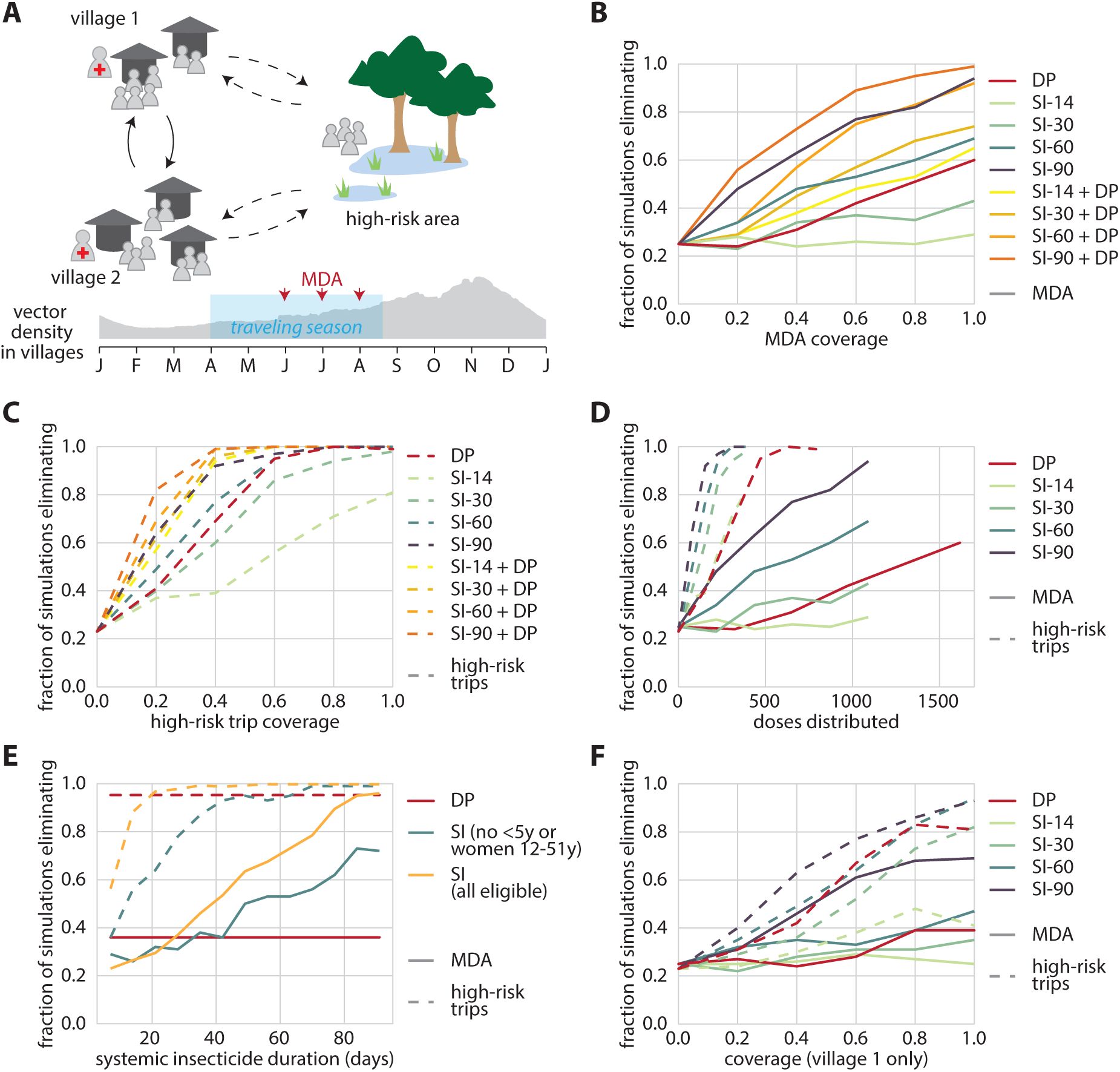
Long-lasting systemic insecticides targeted at mobile, high-risk individuals can lead to elimination. Children under 5 and women of childbearing age are excluded from systemic insecticide eligibility unless otherwise indicated. (A) Model configuration includes two villages and a connected high-risk area to which some villagers regularly travel. Treatment for symptomatic malaria is available in villages but not in the high-risk area. MDA and traveling months occur during the end of the dry season. (B) Likelihood of elimination after MDA in both villages depends on MDA coverage and duration of systemic insecticide activity. (C) Targeting high-risk travelers in both villages as they depart for the high-risk area can be a successful elimination strategy, and minimal coverage required depends on duration of systemic insecticide activity. (D) Targeting high-risk travelers is more efficient at elimination than village-level MDA. (E) Long-lasting systemic insecticides should maintain activity for at least 30-40 days to be more effective than the antimalarial DP at elimination in this context, but improved safety for children and pregnant women would shorten the required systemic insecticide duration. Shown: 60% coverage for both scenarios. (F) Distributing long-lasting systemic insecticides in MDA or to high-risk travelers can still result in likely elimination even if only residents of a single village receive the intervention, but requires higher coverage.

High-risk travelers are 70% of people between ages 15 and 35. Migration happens between April and August with a mean stay duration of 30 days. During each month of high-risk travel, 50% of high-risk travelers who are currently at home in their village can make a trip to the high-risk area. We assume equal gender representation in the high-risk traveler population.

Treatment with artemether-lumefantrine (AL) is available in both villages for symptomatic cases. 80% of clinical malaria cases in children under 10 receive treatment, 70% of clinical cases in individuals over 10 receive treatment, and 95% of severe cases receive treatment. When treatment is sought, it is received within 3 days of symptom onset. High-risk travelers seek care at a lower (40%) and slower (within 5 days) rate. No treatment is available in the high-risk area and no other vector control is simulated.

All simulations last 3 years. We consider distribution of DP only, systemic insecticide only, and DP in combination with systemic insecticide. Children under 5 and women of childbearing age (12-51 years) are ineligible to receive systemic insecticides unless otherwise indicated but can still receive DP. MDA distributions occur only in year 1 and include three independent rounds separated by 30 days, beginning in June. MDA is never distributed in the high-risk area, and coverage refers to the fraction of eligible individuals residing in the village on the date of distribution who receive drugs. In scenarios where individuals receive drugs when they depart the village for the high-risk area, these trip-based drug distributions occur during all years. Coverage refers to the probability that drugs are taken for any given trip to the high-risk area by an eligible individual, and there are no restrictions on the number of times a person can take drugs.

Elimination is defined as zero infected individuals in the entire modeled area, including the high-risk area, and is assessed at the end of the third year. Each scenario is run for 100 stochastic realizations.

### Southern Africa elimination scenarios

We model a 10,000-person population center as 332 separate 1 km^2^ grid cells. Individuals move between grid cells according to a gravity model, with people taking an average of 5 overnight trips per year. There is no disease importation from outside the modelled area. Transmission intensity and seasonality reflect southern Africa, with two vector species: *Anopheles arabiensis* (80% of vectors) and *Anopheles funestus* (20% of vectors) [26,27]. Both vector species are assumed to have 65% anthropophily, with 50% indoor biting for *An. arabiensis* and 90% for *An. funestus.* Parasite prevalence by rapid diagnostic test (RDT) fluctuates between 20% and 40% prior to any vector control or MDA.

When presenting with a clinical episode, 60% of children under 5 and 40% of individuals over 5 receive treatment. 80% of severe clinical episodes are treated for all age groups. Cases that receive care are treated with AL.

Intervention scenarios are simulated for four years (Figure 3A). ITN distributions occur on September 1^st^ of years 1 and 3. ITN killing efficacy starts at 60% and exponentially decays with a time constant of 4 years, and blocking efficacy starts at 90% and exponentially decays with a time constant of 2 years. Individuals age 5 to 20 years old are 10% less likely to use a net [28]. Modeled ITN usage is seasonal, with a maximum of 100% of ITN owners using their net on January 1^st^, reaching its minimum of 50% in June. To capture incomplete net retention, we model 60% of the population discarding their nets at a rate with exponential time constant 260 days and the remaining 40% of individuals keeping their nets for much longer, discarding with an exponential decay time constant of 2160 days [29,30]. Coverage across ITN distributions is uncorrelated.

**Figure 3.**
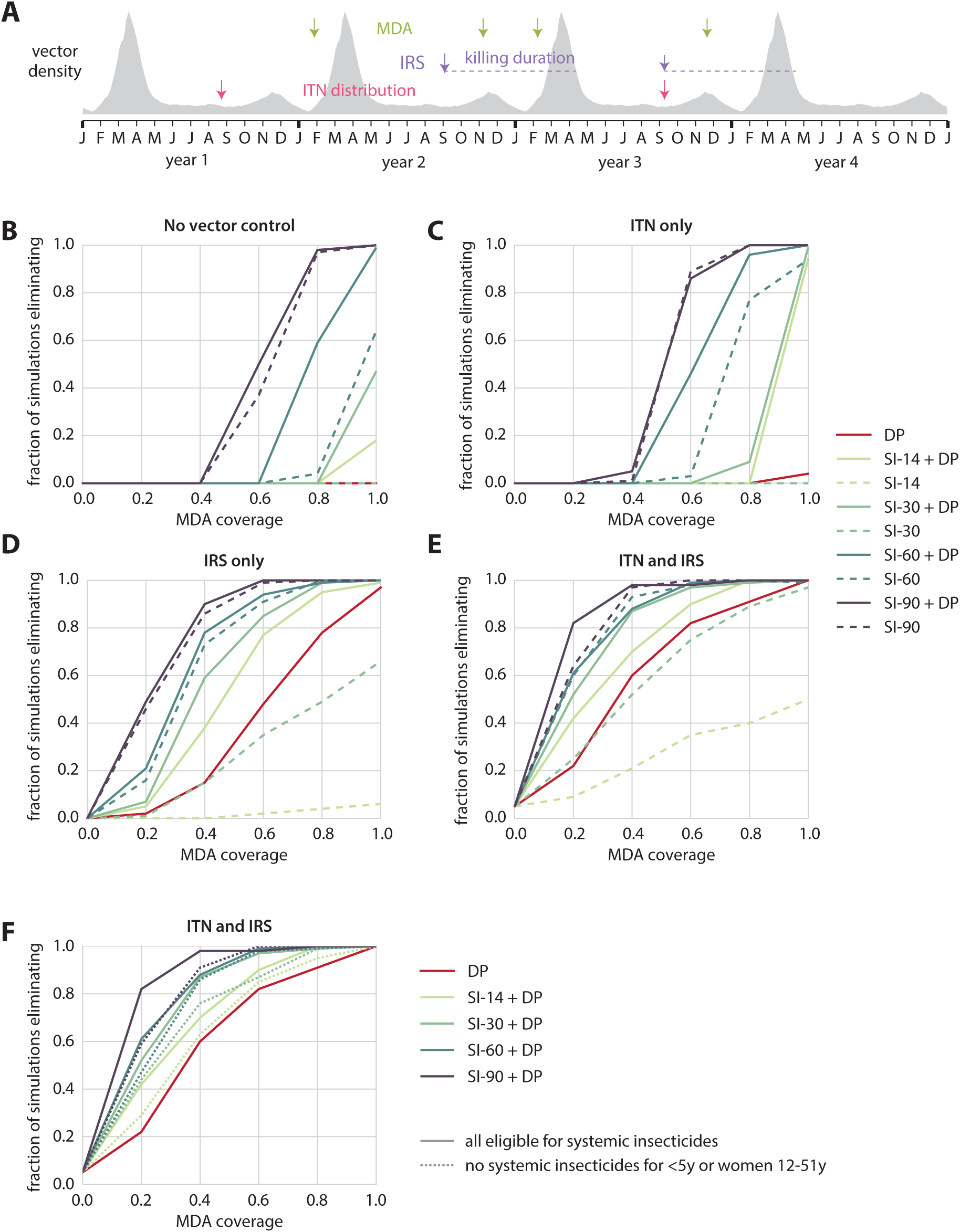
Long-lasting systemic insecticides can complement pre-existing effective vector control in Southern Africa elimination scenarios. (A) Timing of interventions, compared to seasonal vector density profile. Elimination is defined as zero infected individuals at the end of year 4. (B)-(F) Fraction of simulations, out of 100 stochastic realizations, that achieve elimination, for a given vector control and MDA package, with no restrictions on systemic insecticide eligibility. Coverage of ITN and IRS, when implemented, is 60%. (B)-(E): Solid lines: DP included in MDA. Dashed lines: MDA with systemic insecticide alone. (F) Solid lines: no restrictions on systemic insecticide eligibility. Dashed lines: children under 5 and women of childbearing age ineligible for systemic insecticides.

IRS spraying campaigns occur on September 1^st^ of years 2 and 3. We model an organophosphate insecticide with 90% initial killing rate that lasts for 7 months then rapidly decays [31] (and unpublished observations from Mara Maquina). Coverage across IRS campaigns is uncorrelated.

MDA campaigns are implemented in both February and November of years 2 and 3, a total of 4 distributions. We incorporate a small scatter in grid cell-level campaign dates such that each full MDA distribution completes over the course of a few weeks. We test the impact of DP-only MDA, systemic insecticide-only MDA, and MDA with both DP and systemic insecticide. Coverage across multiple MDA rounds is uncorrelated.

No interventions other than ongoing case management are implemented in year 4. Elimination is defined as zero infected individuals in the entire simulated area at the end of year 4. Each intervention scenario is simulated with 100 stochastic realizations.

## Results

### Systemic insecticides lasting at least 60 days reduce burden when given in combination with SMC

The Sahelian setting is characterized by a single peak transmission season and low levels of biting throughout the remainder of the year (Figure 1A). SMC is a commonly used intervention for burden reduction in the Sahel [32], where burden is still high. In a recent study, systemic insecticides were distributed over 6 rounds at 3-week intervals, alongside an SMC campaign, to everyone excluding young children, pregnant women, and women nursing infants under 1 week old, resulting in a 20% drop in clinical incidence [16]. However, short-duration systemic insecticides distributed over multiple rounds may prove operationally challenging at scale, especially during the wet season. Considering this, the existing operational framework of SMC can be leveraged to distribute a single round of long-lasting systemic insecticide with the first round of SMC just as the wet season begins, making it a more tractable intervention configuration for further gains in burden reduction.

We first test the scenario in which all individuals can receive systemic insecticides. We show the proportion of cases remaining after SMC that are averted with systemic insecticide MDA (Figure 1B). As coverage increases, the absolute number of cases available for the systemic insecticide MDA to avert decreases since the SMC campaign is having a greater impact. We find that only SI-60 and SI-90 have meaningful impact on reducing clinical burden. A single round of MDA with systemic insecticides of duration 30 days or less is unlikely to boost the impact of a standard SMC campaign in children under 5. At 60% coverage, SI-90 results in 80% further reduction in burden while SI-30 results in under 10% further reduction in burden compared to a standard SMC campaign at 60% coverage.

If only children are excluded from the MDA target population, coverage as low as 40% with SI-90 is sufficient to begin seeing reduction of the remaining burden by over 20% (Figure 1C). At 60% coverage, simulations predict that SI-60 will also result in around 20% burden reduction, but shorter-lasting systemic insecticides continue to have minimal impact. We observe a 17% reduction in incidence with SI-60 in children under 5 despite this group being excluded from the systemic insecticide MDA (Additional file 1 row B), which is similar to results from the field trial of 6 MDA rounds of ivermectin [16].

We then test the scenario where both children under 5 and women of childbearing age (12 – 51 years) are ineligible to receive long-lasting systemic insecticides (Figure 1D), which describes a case where systemic insecticide safety in young children and pregnant women has not yet been established and pregnancy testing is unavailable during MDA. These populations do not enter into the denominator of coverage calculations for the MDA. Under this restricted distribution scenario, only SI-90 resulted in substantial additional reduction in clinical burden relative to SMC alone, and furthermore MDA coverage of at least 60% was required. Given that the MDA-eligible population in this scenario consists primarily of adult men, achieving high coverage may be particularly operationally challenging [33].

At 60% coverage, adding systemic insecticides to a standard SMC campaign results in a 20% drop in annual EIR with SI-14, and a 40% drop in annual EIR with SI-30 even when women of childbearing age and children under 5 are excluded from the systemic insecticide MDA (Additional file 1). When everyone is eligible to receive systemic insecticides at 60% coverage, annual EIR is reduced by 40% and 60% when using SI-14 and SI-30, respectively. However, despite these large drops in EIR, transmission remains high, and only systemic insecticides of durations 60 days or longer reduce EIR to levels that reflect meaningful impact in terms of reduced clinical burden.

A single round of SI-30 or SI-14, concurrently distributed with a standard SMC campaign, has minimal effect on further reducing burden even at 100% coverage. Given safety concerns surrounding the administration of this class of drugs to women of childbearing age [13] and children under 5 [16], only long-lasting systemic insecticides of duration 90 days improve burden reduction in a meaningful way. However, in the event this class of drugs is found to be safe for use in nursing and pregnant women, and children under 5 years, a single round of SI-60 in parallel with standard SMC can still further reduce burden with coverage of around 60%.

Compared to a standard SMC campaign, expanding SMC to include children up to the age of 10 reduces burden in all campaigns with or without systemic insecticides (Figures 1E-G). Without systemic insecticides, the expanded SMC campaign at 40% coverage reduces burden by 20% over a standard SMC campaign. As in the standard SMC use case, systemic insecticides of duration under 30 days do not offer any substantial gains in burden reduction when administered alongside an expanded SMC campaign.

When children under 5 are excluded from the MDA campaign (Figure 1F), 60% coverage results in 90% reduction in burden with SI-90 while SI-60 requires 100% coverage to offer the same 90% reduction in burden. Excluding children under 5 produces similar results to a campaign where everyone is included (Figures 1E and 1F). Just 50% coverage with SI-90 results in over 80% reduction in burden.

When women of childbearing age and children under 5 are excluded from systemic insecticide distribution, 60% coverage with SI-90 results in 80% reduction in burden, and 100% coverage with SI-60 results in just over 80% reduction in burden (Figure 1G). SI-90 with an expanded SMC campaign, excluding women of childbearing age and children under 5, achieves similar burden reduction as SI-90 with a standard SMC campaign and no restrictions on systemic insecticide eligibility. However, with SI-60 the scenario with expanded SMC eligibility and restricted systemic insecticide eligibility performs better than the scenario with standard SMC and unrestricted systemic insecticide.

Given the high rates of burden experienced in children under the age of 15 [13], expanding an SMC campaign to include children up to the age of 10 could be highly beneficial in many high-transmission settings. If duration of systemic insecticide effectiveness is at least 60 days, long-lasting systemic insecticides further reduce burden when added to an SMC campaign. If malaria control programs have the capacity to distribute multiple rounds of systemic insecticides to older children and adults alongside an SMC campaign, shorter duration systemic insecticides could still be a worthwhile option [13]. Ultimately, safety and cost-effectiveness of the drugs used in systemic insecticide MDA will determine which campaign – expanded or standard SMC with regular or long-lasting systemic insecticides – is most efficacious in reducing burden for a given setting.

### Targeting high-risk travelers with long-lasting systemic insecticides can be part of an effective elimination strategy

To explore the potential role of long-lasting systemic insecticides in reaching elimination in low-transmission settings with mobile populations, we simulate a system of two villages connected to each other and to a shared high-risk area, which is visited by a subset of villagers (Figure 2A). Case management for uncomplicated and severe malaria is available in the villages but not in the high-risk area. Transmission in the entire system is low enough that treatment-seeking alone results in interruption of malaria transmission within 3 years in 25% of the simulations.

We test two modes of systemic insecticide delivery: (1) as MDA distributed in the villages and (2) given to high-risk travelers as they depart for the high-risk area. We assume that children under 5 and women of childbearing age are ineligible to receive systemic insecticide but can still receive DP. In the MDA scenario, achieving elimination with at least 80% probability is possible at operationally feasible coverage with SI-60 when co-administered with DP, or SI-90 if systemic insecticide is given without an antimalarial (Figure 2B). MDA with SI-90 requires 70% coverage to achieve 80% probability of elimination without co-administration of DP, or 50% coverage if the MDA also includes DP. Systemic insecticides of shorter duration or DP-only MDA fail to reach 80% probability of elimination even at 100% coverage.

DP-only MDA is as or more successful at elimination at all coverage levels compared with systemic insecticide MDA when systemic insecticide duration is 30 or fewer days. If DP-only MDA is already planned, addition of SI-14 increases probability of elimination by only a small amount. If MDA with systemic insecticide is already planned, addition of DP to the MDA regimen is highly beneficial for all systemic insecticide durations tested, although smaller gains are seen with SI-90, which is already highly successful. If safety concerns in children and pregnant women are alleviated, the combination of DP with SI-30 leads to elimination with 80% probability when coverage is above 80% (Additional File 2).

Compared with MDA, eliminating malaria with a traveler-targeted drug distribution requires lower coverage (Figure 2C). When systemic insecticide lasts for at least 60 days, coverage as low as 45% results in highly likely elimination, and even DP or SI-30 can eliminate with coverage below 60%. There is some additional gain in performance if systemic insecticide duration is increased above 60 days, as SI-90 requires only 30% coverage to achieve likely elimination. Only 30% coverage is required for likely elimination if travelers are given both DP and systemic insecticide, even if the systemic insecticides lasts only 14 days. The model assumes men and women are equally likely to be high-risk travelers, and thus only half the high-risk trips are eligible to receive systemic insecticide. If this assumption is relaxed, coverage as low as 30% with SI-30 can result in >80% probability of elimination in the trip-based distribution scenario (Additional File 2). The traveler-targeted strategy requires fewer doses of drug than MDA to achieve the same level of success (Figure 2D), even though the traveler-targeted strategy is implemented for 3 years while MDA is implemented for one.

Fixing coverage at 60%, we sampled systemic insecticide durations from 7 days to 98 days at 1-week increments and compared the likelihood of elimination against the DP alternative **(Error! Reference source not found. 2E)**. To outperform DP, systemic insecticide should last at least 40 days when given as MDA or 50 days if given to high-risk travelers. If systemic insecticide can be given to children and women of childbearing age, the minimum duration for systemic insecticide to outperform DP as an elimination drug is reduced to 30 days for MDA and only 20 days for the trip-based distribution.

If only one of the two villages can be reached by either MDA or the trip-based intervention, elimination is still possible, but higher coverage is required (Figure 2F). Human movement connects populations and is often discussed as a barrier to achieving malaria elimination. In this context, however, when transmission is sustained by movement to a high-risk area where other types of interventions are not easily applied, humans carrying long-acting drugs in their bodies could provide an effective method for delivering vector control to hard-to-reach areas.

### Systemic insecticides decrease the MDA campaign coverage needed to interrupt transmission in an integrated intervention strategy

To explore the role of long-lasting systemic insecticides in an elimination strategy that includes traditional vector control and MDA with antimalarials, we simulate a spatially distributed area with southern-Africa-like transmission dynamics (Figure 3A). To isolate the intervention effects, we assume no importation of infections into this area. Since *An. arabiensis* comprises about 80% of the vector density, there is a substantial amount of outdoor biting.

Figure 3 shows the likelihood of elimination under four possible underlying vector control packages: no traditional vector control, ITNs only, IRS only, and ITN+IRS. Under no traditional vector control, an MDA campaign with systemic insecticide is highly unlikely to achieve elimination, unless the systemic insecticide is very long-lasting and coverage is very high: over 70% coverage for SI-90-only MDA or 90% coverage for MDA with SI-60 + DP (Figure 3B). If the underlying vector control is ITNs alone, the situation is largely the same (Figure 3C); however, if the underlying vector control is IRS alone, elimination is much more likely (Figure 3D). This difference is due to two effects: first, we have assumed significant pyrethroid resistance limiting the ITN killing efficacy and little resistance to the organophosphate used in IRS. The second effect is that half the modeled population discards their ITNs within 18 months, whereas the IRS spray, not subject to behavioral factors, is assumed stable for the full 7 months of its duration. In the context of IRS without ITNs, systemic insecticide duration can compensate for MDA coverage (Figure 3D). To achieve elimination with 80% probability, a DP-only MDA requires 85% coverage, an MDA with DP and SI-14 requires 65% coverage, and an MDA with DP and SI-90 requires only 35% coverage. If the vector control package includes both ITN and IRS, the qualitative picture is similar to the IRS-only case but elimination becomes more feasible across the board, further reducing the MDA coverages needed (Figure 3E).

When vector control is poor, a systemic insecticide only adds additional benefit to an already-planned MDA with DP if the systemic insecticide duration is greater than 60 days (Figure 3B, 3C). In the case of a DP + systemic insecticide MDA drug package combined with a vector control program including IRS, however, the stronger vector control foundation enables systemic insecticides of any duration to give substantial added benefit alongside DP. Focusing on SI-14, whose duration of activity is close to that of high-dose ivermectin, we find that SI-14 adds value to a DP-only MDA only when IRS is implemented and when vector control coverage is good but not excellent, above 40% but under 80% (Figure 4). Below this level of vector control coverage, elimination is highly unlikely and the SI-14 does not have much impact; above this coverage, DP alone is sufficient to make elimination likely. In this intermediate regime of vector control coverage, where many areas that have implemented IRS are likely to lie, adding SI-14 to DP increases the chance of elimination by 10-40%. The synergy between IRS and MDA with antimalarials has previously been described [34], and we find further opportunity to boost this interaction by incorporating systemic insecticide in the MDA.

**Figure 4.**
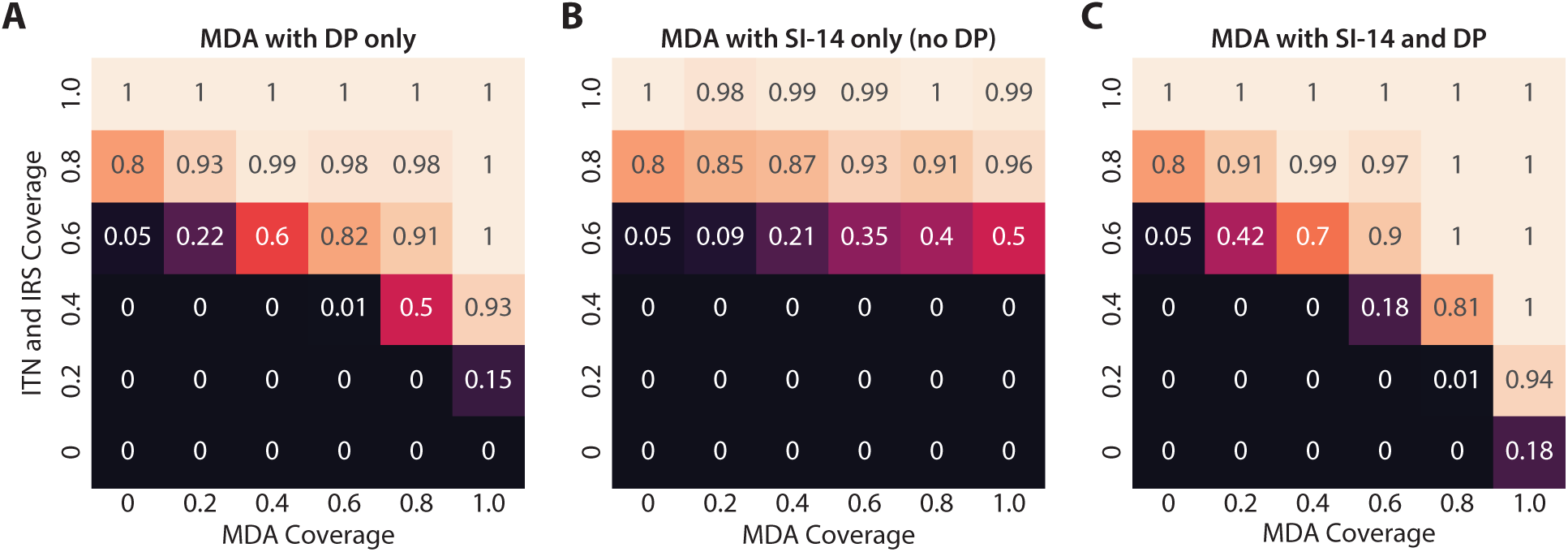
Southern Africa elimination outcomes for MDA packages with (A) DP only (B) SI-14 only or (C) both DP and SI-14. A systemic insecticide-only MDA with 14-day systemic insecticide duration is always inferior to a DP-only MDA at the same coverage. However, an SI-14+DP MDA is superior to a DP-only MDA, for operationally realistic vector control and MDA campaign coverages. The number in each cell is the fraction of simulations, out of 100 stochastic realizations, that achieved elimination for the corresponding vector control and MDA coverages. Each row of the figure corresponds to a different vector control scenario, and each column of the figure corresponds to a different MDA drug package.

We also investigate the alternative use case where a systemic insecticide MDA is already planned and determine when adding DP to the MDA adds value. When systemic insecticide is included in an MDA, the addition of DP typically has the strongest effect for SI-14 and SI-30 but has less marginal benefit for longer-lasting systemic insecticides. Figure 4, which focuses exclusively on the impact of SI-14, shows that at any coverage of traditional vector control coverage, adding DP to a 14-day systemic insecticide always increases the probability of elimination. For SI-90, there is minimal benefit from adding DP, while for SI-60, adding DP to the MDA regimen is substantially beneficial only if MDA coverage is high and traditional vector control is poor.

Unlike the SMC and targeted elimination use cases, the Southern Africa scenarios we have investigated thus far allowed children under the age of 5 and women of childbearing age to receive systemic insecticide. Restricting these populations’ eligibility for systemic insecticide predictably requires higher campaign coverages to have the same effect, by roughly 5-15%, depending on the systemic insecticide duration (Figure 3F). For SI-14, a restricted systemic insecticide distribution reduces its effectiveness to the point that the systemic insecticide provides very little benefit above a DP-only campaign. For longer-lasting systemic insecticides, even a restricted systemic insecticide + DP campaign has larger impact than a DP-only campaign.

Robust vector control is a crucial foundation towards achieving elimination in the Southern Africa context. Long-lasting systemic insecticides complement but do not replace traditional vector control, and nor do they replace antimalarials during MDA unless the systemic insecticide duration is 90 days. However, including systemic insecticides in an antimalarial MDA can substantially lower the coverage required to reach elimination, which could be operationally attractive since absence and exclusion criteria can limit the maximum achievable coverage.

## Discussion

In this work, we explore the efficacy of systemic insecticides as a malaria control tool in three different transmission settings: a Sahelian setting with high annual EIR where the goal is burden reduction, a near-elimination setting with a group of adults who regularly visit high-risk areas, and a southern Africa context focusing on elimination as the desired endpoint through vector control and drug campaigns. These diverse settings were selected to identify opportunities where systemic insecticide could provide high value to a pre-existing drug delivery program, and to explore some of the very different contexts in which additional malaria control tools might help accelerate burden reduction or elimination.

Across the three modeled settings, the use case for including a systemic insecticide is strongest when the systemic insecticide duration is greater than about 30 days in a near-elimination setting, or greater than 60 days in a high-burden setting. Distributing systemic insecticide that lasts only 14 days had little to no impact in all of the modeled settings. In a near-elimination context with robust vector control, good campaign coverage, and, importantly, a safety profile sufficient to be given to small children and women of childbearing age, a 14-day systemic insecticide can be beneficial.

We predict that systemic insecticides offer the greatest benefit when layered on top of other interventions rather than administered in isolation. In the southern Africa context, elimination is highly unlikely without a foundation of robust vector control, and systemic insecticides add the most benefit when administered alongside an integrated system of vector control and DP drug campaigns; here, the systemic insecticide can substantially reduce the MDA campaign coverage needed to achieve elimination. Lowering the MDA coverage necessary to achieve elimination is an important operational consideration, partly because mobile, high-risk groups are often difficult to reach. However, if these high-risk groups can be targeted directly, for example by treating adults in Southeast Asian settings who enter the deep forest for work with systemic insecticide and DP before they set out, the interventions have an outsize impact on transmission. When close to elimination, trying to target the sources of imported cases may be more cost-effective than continuing to apply interventions in population centers.

Children under the age of 5 and women of childbearing age comprise about 38% of the population in SE Asia and 44% of the population in Sub-Saharan Africa. A systemic insecticide that cannot be safely given to these large portions of the population is more limited in its potential impact, although we have identified cases where such a restricted systemic insecticide may still be still beneficial. We have assumed no pregnancy testing during MDA in order to minimize complexity and cost of MDA programs, although if pregnancy testing is an option, many more people would be able to take the systemic insecticide. If a systemic insecticide is unsafe to give to vulnerable subgroups, then to have the same efficacy, either the campaign coverage or the systemic insecticide duration must be increased. For example, in the high-transmission Sahelian context, an unrestricted systemic insecticide of 60-day duration has a similar impact on clinical burden as a restricted systemic insecticide of 90-day duration. In the targeted elimination scenario, systemic insecticide safety restrictions can more than double the systemic insecticide duration necessary to achieve elimination. Prior to approval for new long-lasting systemic insecticides in children and pregnant women, these groups could instead receive regular dose ivermectin [35].

We simplify the pharmacokinetic and pharmacodynamic properties of systemic insecticides by assuming a constant killing efficacy over the course of the insecticide’s duration, allowing us to isolate the impact of insecticide duration on epidemiological outcomes. The implications for more complex within-host insecticide dynamics can be interpolated from the results presented here. For example, a 300ug/kg ivermectin dose can be compared with the modeled hypothetical SI-14 [15]. Our model also does not incorporate possible non-fatal vector outcomes of the systemic insecticide such as reduced fecundity, nor did we model any parasite killing effects since these outcomes are not well understood and the effects are likely secondary to killing the mosquito [19].

The scenarios presented in this work do not directly incorporate effects of drug or insecticide resistance, the exception being in assuming a low killing efficacy of ITNs in the Southern Africa scenario due to pyrethroid resistance, which is widespread [36]. In settings where resistance is present, systemic insecticides of a shorter duration than 30 or 60 days could still be an important operational tool. Against the backdrop of growing insecticide resistance, systemic insecticides have added utility since they operate by a novel mechanism not shared by current ITN and IRS insecticide compounds, and there is potential for synergy between the insecticides’ killing mechanisms. Against drug resistance, systemic insecticides layered on top of an antimalarial could also be beneficial, since even if parasites have some protection against the antimalarial, the systemic insecticide may be able to kill most of the vectors carrying this resistant strain. If systemic insecticide replaces DP entirely for mass administrations, selection pressure could be partially transferred from the parasite onto the vector, making further development of antimalarial drug resistance less likely.

Anthropophily, the fraction of bites that are taken on a human host, is a critical vector bionomic parameter that is often poorly understood in the local vector context. For a fixed mosquito population size, lowering anthropophily lowers the number of bites on humans and thus reduces the chance of a mosquito transmitting malaria. However, a lower anthropophily also makes the mosquito population more resilient to the impact of vector control methods such as ITN, IRS, or systemic insecticide where vectors encounter vector control while seeking human hosts. We find that longer-lasting systemic insecticides are less sensitive to uncertainty in anthropophily than DP or systemic insecticides of duration 30 days or less, and that probability of elimination decreases as anthropophily increases regardless of the drug deployed (Additional File 2). We did not consider distribution of systemic insecticides to livestock, although this strategy may be fruitful where vectors are fairly zoophilic and humans and livestock live in close proximity [37].

While we find that long-lasting systemic insecticides can yield large benefits in terms of burden reduction or hitting elimination targets, given a good safety profile, co-administration with antimalarials, and a foundation of robust traditional vector control, the operational challenges and costs of administering MDA should not be ignored when considering this intervention. Other new tools for control of outdoor-biting vectors are also under development, and the cost-effectiveness of systemic insecticides should be compared with other available options.

## Conclusion

When distributed in a single round alongside SMC, only systemic insecticides of duration 60 or more days have a meaningful impact on reducing the overall clinical burden. In a near-elimination setting with high-risk travelers, targeting long-lasting systemic insecticides to seasonal high-risk travelers can be a highly effective elimination strategy, more effective than MDA in the general population. In a southern Africa elimination setting that includes existing IRS, ITN, and MDA campaigns with antimalarials, adding systemic insecticide to the MDA regimen increases the probability of elimination and lowers the MDA coverage required to achieve elimination. Systemic insecticides lasting 40 days or more are more effective than the antimalarial DP for elimination. However, systemic insecticides of durations between 14 and 30 days can still be effective tools in lower-transmission settings if young children and pregnant and nursing women can be treated safely.

## List of abbreviations

AL: artemether-lumefantrine
DP: dihydroartemisinin-piperaquine
EIR: entomological inoculation rate
SI-14: systemic insecticide with 14-day duration
SI-30: systemic insecticide with 30-day duration
SI-60: systemic insecticide with 60-day duration
SI-90: systemic insecticide with 90-day duration
IRS: indoor residual spray
ITN: insecticide-treated net
MDA: mass drug administration
SMC: seasonal malaria chemoprevention

## Declarations

### Ethics approval and consent to participate

Not applicable

### Consent for publication

Not applicable

### Availability of data and material

All software used to generate and analyze the data in this manuscript are publicly available or available upon request as described in Additional File 3.

### Competing interests

The authors declare that they have no competing interests.

## Funding

This work was supported by Bill and Melinda Gates through the Global Good Fund.

## Authors’ contributions

JG and EW conceived the project. PS, JS, and JG designed, built, and ran the simulations. PS, JS, CB, and JG analyzed the data and wrote the manuscript. All authors read and approved the final manuscript.

### Acknowledgements

The authors thank Svetlana Titova for software support and Carlos Chaccour, Janice Culpepper, and Erin Stuckey for helpful discussions.

## Additional Files

Additional File 1. Impact of long-lasting systemic insecticides combined with standard SMC on annual EIR and case reduction by age group.

Additional File 2. Impact of long-lasting systemic insecticides in a targeted elimination scenario where children under 5 and women of childbearing age are eligible to receive systemic insecticide.

Additional File 3. Directions on how to obtain the software and scripts used to generate and analyze the data in this study.

