## Supplementary figures and images for "Reducing malaria burden and accelerating elimination with long-lasting systemic insecticides: a modeling study of three potential use cases"

### Additional File 1

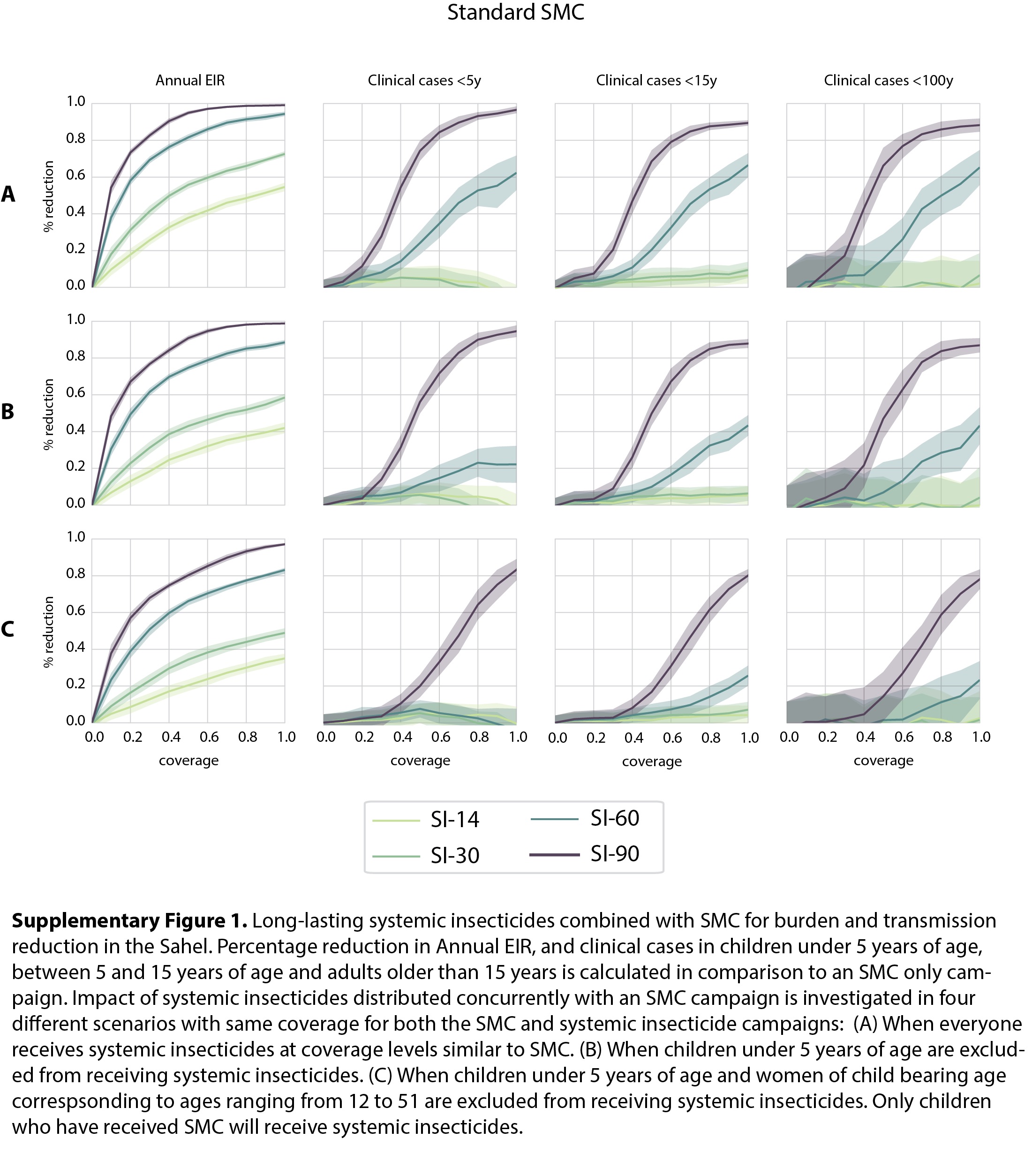

### Additional File 2

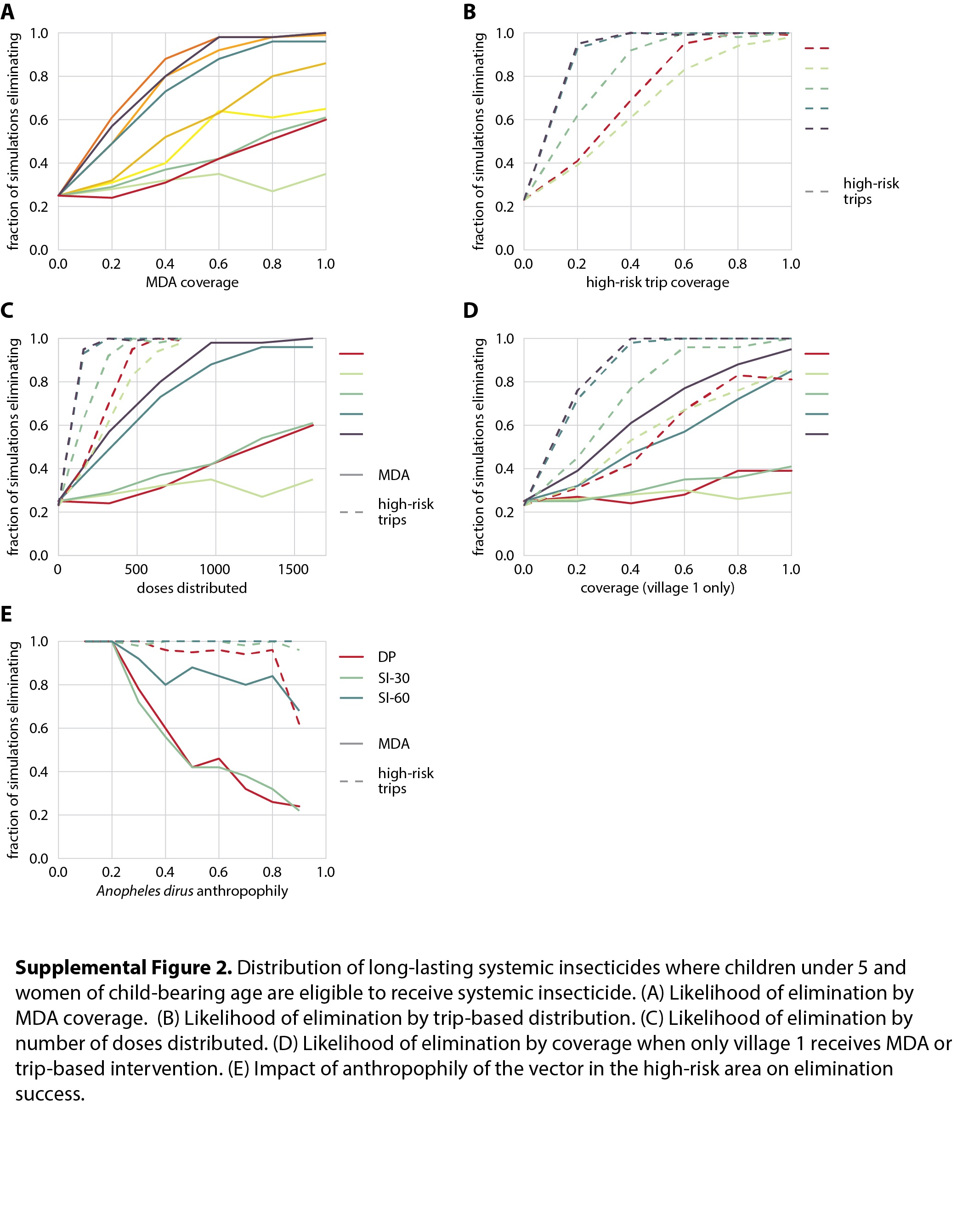
